# Bidirectional Wnt signaling between endoderm and mesoderm confer tracheal identity in mouse and human

**DOI:** 10.1101/758235

**Authors:** Keishi Kishimoto, Kana T. Furukawa, Agustin Luz Madrigal, Akira Yamaoka, Chisa Matsuoka, Masanobu Habu, Cantas Alev, Aaron M. Zorn, Mitsuru Morimoto

**Affiliations:** Laboratory for Lung Development, Riken Center for Biosystems Dynamics Research (BDR), Kobe, 650-0047, Japan; RIKEN BDR – CuSTOM Joint Laboratory, Cincinnati Children’s Hospital Medical Center, Cincinnati, OH 45229, USA; Center for Stem Cell & Organoid Medicine (CuSTOM), Cincinnati Children’s Hospital Medical Center, Cincinnati, OH 45229, USA; Division of Developmental Biology, Cincinnati Children’s Hospital Medical Center, Cincinnati, OH 45229, USA; Department of Cell Growth and Differentiation, Center for iPS Cell Research and Application (CiRA), Kyoto University, Kyoto 606-8507, Japan; Institute for the Advanced Study of Human Biology (ASHBi), Kyoto University, Kyoto 606-8501, Japan

## Abstract

The periodic cartilage and smooth muscle structures in mammalian trachea are derived from tracheal mesoderm, and tracheal malformations result in serious respiratory defects in neonates. Here we show that canonical Wnt signaling in mesoderm is critical to confer trachea mesenchymal identity in human and mouse. Loss of β-catenin in fetal mouse mesoderm caused loss of Tbx4^+^ tracheal mesoderm and tracheal cartilage agenesis. The Tbx4 expression relied on endodermal Wnt activity and its downstream Wnt ligand but independent of known Nkx2.1-mediated respiratory development, suggesting that bidirectional Wnt signaling between endoderm and mesoderm promotes trachea development. Repopulating *in vivo* model, activating Wnt, Bmp signaling in mouse embryonic stem cell (ESC)-derived lateral plate mesoderm (LPM) generated tracheal mesoderm containing chondrocytes and smooth muscle cells. For human ESC-derived LPM, SHH activation was required along with Wnt to generate proper tracheal mesoderm. Together, these findings may contribute to developing applications for human tracheal tissue repair.

## Main paragraph

The mammalian respiratory system is crucial for postnatal survival, and defects in the development of the respiratory system cause life-threatening defects in breathing at birth^1^. The trachea is a large tubular air path that delivers external air to the lung. Abnormal development of the tracheal mesenchyme, including cartilage and smooth muscle (SM), is associated with congenital defects in cartilage and SM such as tracheoesophageal fistula (TEF) and tracheal agenesis (TA)^2, 3^. Thus, understanding trachea development is crucial to better understand TEF/TA and establish a protocol to reconstruct trachea from pluripotent stem cells for human tissue repair.

Trachea/lung organogenesis is coordinated by endodermal-mesodermal interactions during embryogenesis. The primordial tracheal/lung endoderm appears at the ventral side of the anterior foregut at embryonic day 9 to 9.5 (E9.0–9.5) in mouse (Fig. 1a). Previous studies have revealed that development of tracheal/lung endoderm is initiated by graduated expression of mesodermal Wnt2/2b and Bmp4 expression along the dorsal-ventral axis^4-7^. This mesodermal-to-endodermal Wnt and Bmp signaling drives expression of Nkx2.1, the key transcription factor of tracheal/lung lineages^8^, at the ventral side of the anterior foregut, which in turn suppresses Sox2 to segregate these Nkx2.1^+^ endodermal cells from the esophageal lineage. The Nkx2.1^+^ endoderm then invaginates into the ventral mesoderm to form the primordial trachea and lung buds. At the same time, the Sox2^+^ endoderm at the dorsal side develops into the esophagus by E10.5 (Fig. 1a). By recapitulating developmental processes *in vitro*, trachea/lung endodermal cells and differentiated epithelial populations have been generated from both mouse and human pluripotent stem cells^9-11^, and can also be used for disease modeling^12-14^. However, an established protocol for inducing tracheal/lung mesoderm and differentiated mesenchymal tissue from pluripotent cells has not yet been reported because developmental signaling pathways coordinating the mesodermal development are still undefined.

**Figure 1.**
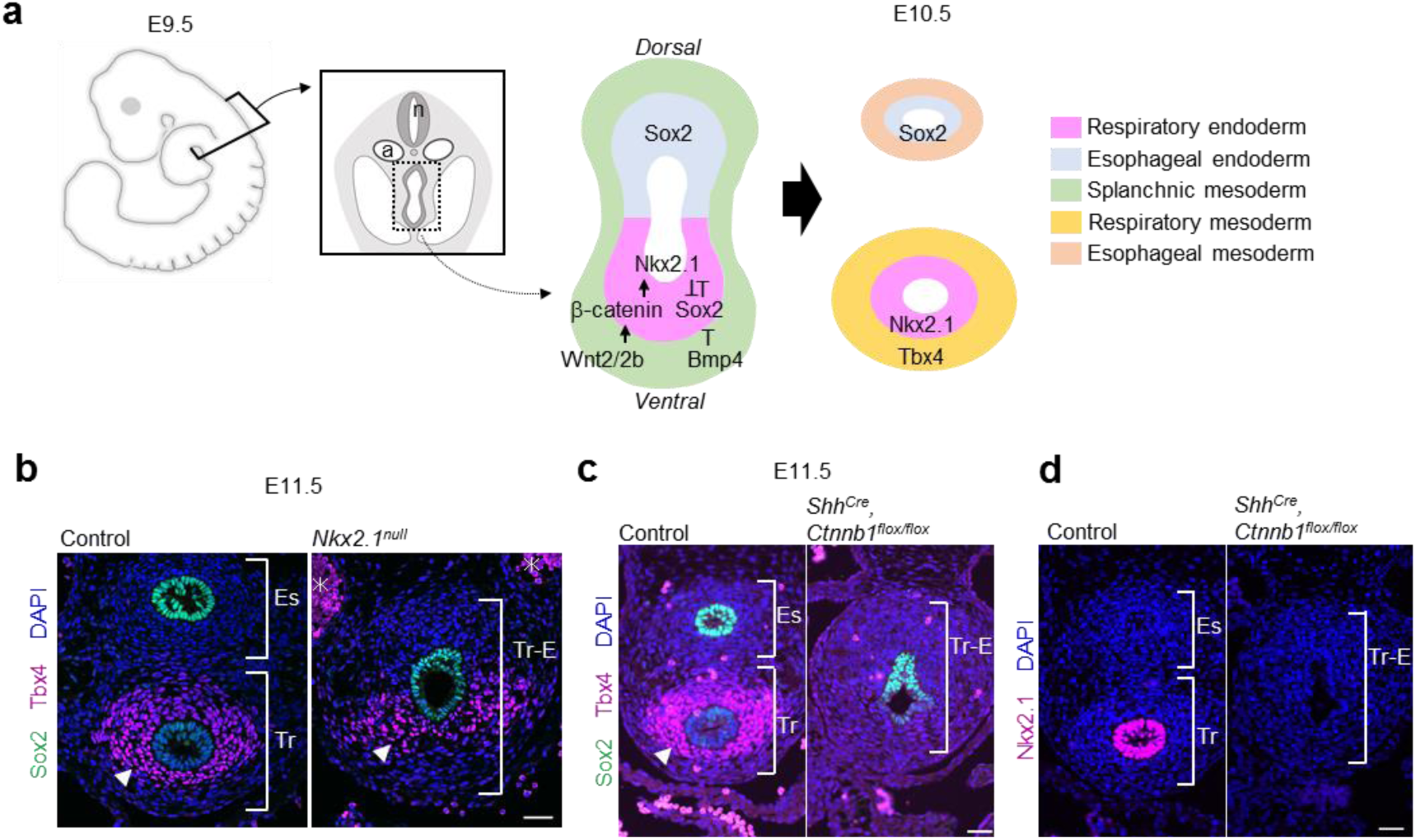
Activation of Wnt signaling in endoderm, but not Nkx2.1 expression, is activated to promote mesodermal development of the mouse trachea. **a**, Schematic model of tracheoesophageal segregation. **b**, Transverse sections of *Nkx2.1*^*null*^ mouse embryos and littermate controls. Sections were stained for Sox2 (*green*), Tbx4 (*magenta*), and DAPI (*blue*). Arrowheads indicate Tbx4^+^ tracheal mesoderm. Asterisks indicate non-specific background signal of blood cells in dorsal aorta. **c**, Transverse sections of *Shh*^*Cre*^, *Ctnnb1*^*flox/flox*^ mouse embryos and littermate controls. Sections were stained by Sox2 (*green*), Tbx4 (*magenta*), and DAPI (*blue*). Arrowheads indicate Tbx4^+^ tracheal mesoderm. **d**, Transverse sections of *Shh*^*Cre*^, *Ctnnb1*^*flox/flox*^ mouse embryos and littermate controls. Sections were stained by Nkx2.1 (*magenta*) and DAPI (*blue*). n; neural tube, a; aorta, Es; Esophagus, Tr; Trachea, Tr-E; Trachea-Esophageal tube Scale bar, 40μm

The tracheal mesoderm originates from ventral fold of the lateral plate mesoderm (LPM) surrounding the anterior foregut endoderm. At the same time as endodermal Nkx2.1 induction, tracheal/lung mesoderm is also defined, expressing Tbx4/5 by E10.5, which are markers for tracheal/lung mesoderm and required for proper mesenchymal development (Fig. 1a)^15^. In contrast to Tbx5 which is also expressed in LPM and cardiac mesoderm^16, 17^, Tbx4 expression is restricted to respiratory tissue. At E9.5, Tbx4 is only detected in lung buds but not tracheal mesoderm (Supplementary Fig. S1). Tbx4 expression is then detected in tracheal mesoderm from E10.5. Tbx4 and Tbx5 cooperate to steer normal trachea development. Both genes are required for mesodermal development of the trachea, particularly for cartilage and smooth muscle differentiation as well as morphogenesis. The crucial functions of these genes are validated by Tbx4, 5 double mutants exhibiting the phenotypes of tracheal stenosis^15^. We previously reported that synchronized polarization of mesodermal cells and temporal initiation of cartilage development regulates tracheal tube morphogenesis by coordinating the length and diameter of the mouse trachea, respectively^18, 19^. However, the mechanism underlying the initial induction of tracheal mesoderm is still unclear.

To study the initiation of the mesodermal development of the trachea, we validated the involvement of Nkx2.1 in mesodermal Tbx4 expression because endodermal-mesodermal interactions orchestrate organogenesis throughout development in general. Nkx2.1 is an endodermal transcription factor necessary for tracheal and lung development and its genetic ablation results in TEF^8^. We examined *Nkx2.1*^*null*^ mouse embryos and confirmed the TEF phenotype with a single tracheoesophageal (Tr-E) tube (Fig. 1b). Interestingly, *Nkx2.1*^*null*^ embryos retained Tbx4 expression in the ventrolateral mesoderm of a single Tr-E tube, although the segregation was defective (Fig. 1b), indicating that mesodermal induction of the trachea is independent of endodermal Nkx2.1. We compared the phenotype of *Nkx2.1*^*null*^ with that of *Shh*^*Cre*^, *Ctnnb1*^*flox/flox*^ embryos which also show anterior foregut endoderm segregation defect and loss of Nkx2.1 expression (Fig. 1c and d)^4, 5^. In contrast to *Nkx2.1*^*null*^ embryos, *Shh*^*Cre*^, *Ctnnb1*^*flox/flox*^ embryos did not express Tbx4, suggesting the activation of endodermal Wnt signaling, but not Nkx2.1, is required for following mesodermal Tbx4 expression. Thus, the initial induction of tracheal mesoderm is independent of known Nkx2.1-mediated respiratory endoderm development, but dependent on Wnt signaling at the ventral anterior foregut endoderm.

To further study the spatiotemporal regulation of canonical Wnt signaling during tracheoesophageal segregation at E9.5 to E11.5, we used a reporter line LEF1 ^EGFP^ and examined the distribution of EGFP in the canonical Wnt signaling response (Figs. 2a and b)^20^. At E9.5, EGFP was detected in the ventral half of the anterior foregut endoderm where trachea endodermal cells appear and express Nkx2.1 (Figs. 2a and b, *arrowheads*) and then decreased temporally at E10.5. After E10.5, the EGFP reporter was activated in the surrounding mesoderm and its intensity increased at E11.5 (Figs. 2a and b, *arrowheads*), which was similar to the patterning of Axin2-LacZ, another reporter line for the response of canonical Wnt signaling^21^. We further conducted RNAscope *in situ* hybridization against Axin2, an endogenous Wnt target gene, to confirm activation of Wnt signaling in mesoderm. Axin2 was highly expressed in surrounding mesoderm at E10.5 compared to endoderm, similar to the pattern observed in the reporter line (Fig. 2c). Because these Wnt-responsive mesodermal cells expressed Tbx4 (Fig. 2b), we hypothesized that Wnt signaling in the early mesoderm is involved in the initiation of the tracheal mesoderm.

**Figure 2.**
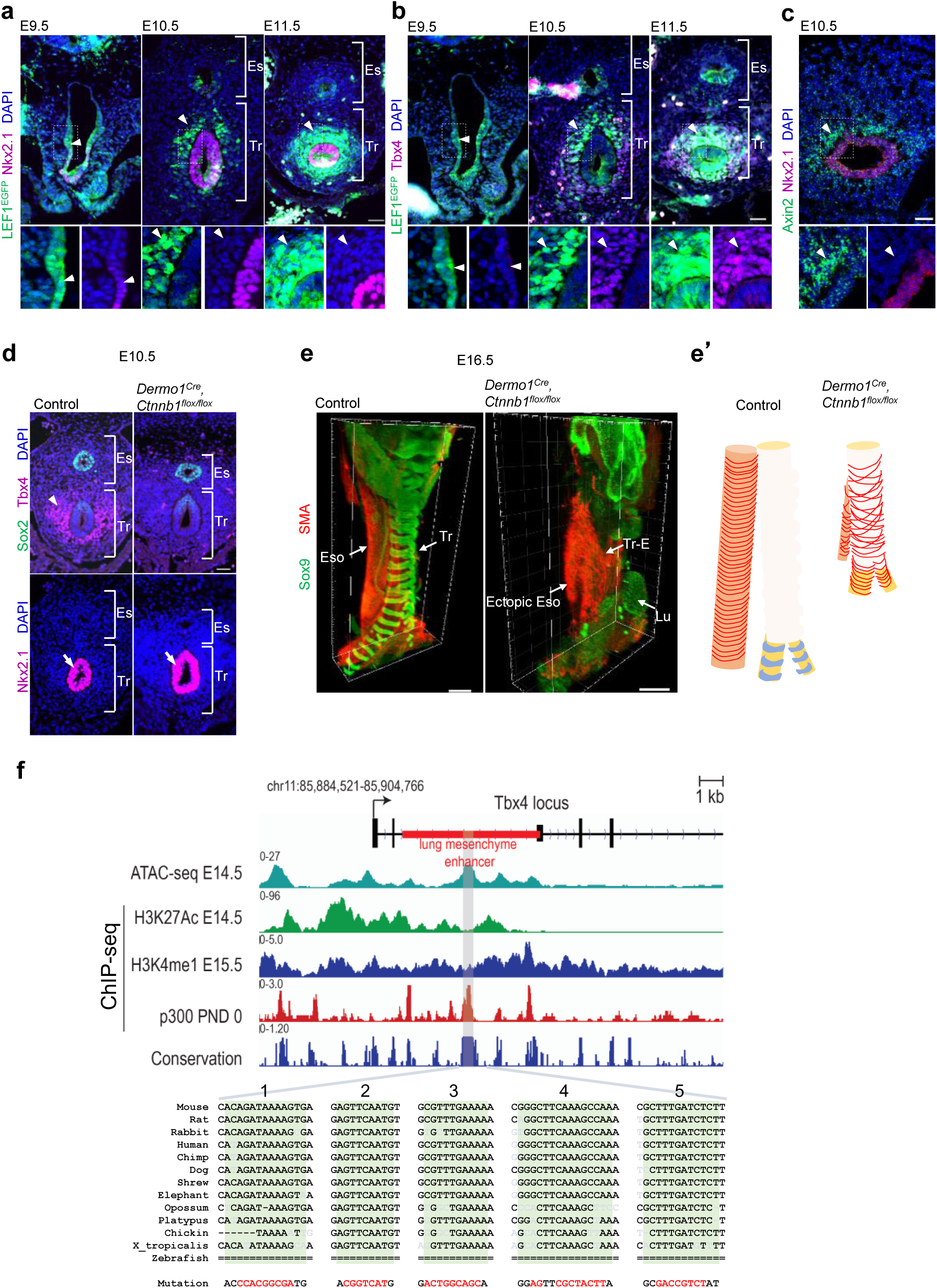
Wnt signaling is activated to promote mesodermal development of the mouse trachea. **a**, Transverse sections of LEF1^EGFP^ reporter mouse embryos at E9.5 to E11.5. Sections were stained for EGFP (*green*), Nkx2.1 (*magenta*), and DAPI (*blue*). Arrowheads indicate GFP^+^ cells. **b**, Transverse sections of LEF1^EGFP^ reporter mouse embryos at E9.5 to E11.5. Sections were stained for EGFP (*green*), Tbx4 (*magenta*), and DAPI (*blue*). Arrowheads indicate GFP^+^ cells. **c**, Transversal section of mouse embryo at E10.5. Section were stained for Axin2 (*green*), Nkx2.1 (*magenta*) and DAPI (*blue*) by RNAscope experiment. Arrowheads indicate Axin2^+^ cells. **d**, Transverse sections of *Dermo1*^*Cre*^, *Ctnnb1*^*flox/flox*^ mouse embryos and littermate controls at E10.5. Upper panels show sections stained for Sox2 (*green*), Tbx4 (*magenta*), and DAPI (*blue*). Lower panels show sections stained for Nkx2.1 (*magenta*) and DAPI (*blue*). Arrowhead indicates Tbx4^+^ cells. Arrows indicate Nkx2.1^+^ cells. **e**, Three-dimensional imaging of whole trachea and esophagus tissue at E16.5. Cartilage morphology and smooth muscle architecture in the tracheas of *Dermo1*^*Cre*^, *Ctnnb1*^*flox/flox*^ mouse embryos and control littermates. Whole trachea and esophagus were stained for Sox9 (*green*) and SMA (*red*). **e’**, Model of tracheal architecture in *Dermo1*^*Cre*^, *Ctnnb1*^*flox/flox*^ mouse embryos and control littermates based on (**e**). Eso; Esophagus, **f**, Integrative Genomics Viewer (IGV) snapshot of mm10 (chr11:85,884,521-85,904,766) showing mouse *tbx4* lung mesenchyme specific enhancer (LME) and compiled ENCODE data of ATAC-seq E14.5 (ENCSR335VJW), H3K27Ac E14.5 (ENCSR452WYC), H3K4me1 E15.5 (ENCFF283EBS), EP300 post-natal day (PND) 0 and vertebrate conservation (Phastcons). Numbers indicate fold enrichment over input (ChIP-seq). CisBP and Jaspar predicted Tcf/Lef-binding sites (highlighted in green, region: mm10, chr11:85,893,703-85,894,206) are localized at the ATAC-seq and p300 peaks that are conserved among most vertebrates. Sequence in red shows the Tcf/Lef-binding sites mutated. Lu; Lung, Tr; Trachea, Tr-E; Tracheoesophageal tube. Scale bar: 40 μm (a, b), 50 μm (c, d), 300 μm (e).

To validate the role of mesodermal Wnt signaling, we genetically ablated Ctnnb1, also known as β-Catenin, which is a core component of canonical Wnt signaling, from embryonic mesoderm. We employed the Dermo1-Cre line which targets embryonic mesoderm, including tracheal/lung mesoderm, and generated *Dermo1*^*Cre*^, *Ctnnb1*^*flox/flox*^ mice^22-25^. In the mutant embryos, Tbx4 expression was absent in the mesoderm at E10.5 (Fig. 2d), indicating that mesodermal canonical Wnt signaling is necessary for Tbx4 expression. In contrast, endodermal Nkx2.1 expression and tracheoesophageal segregation were not affected, implying that mesodermal Wnt signaling and Tbx4 is dispensable for endodermal development. Supporting the observation of lung buds in *Dermo1*^*Cre*^, *Ctnnb1*^*flox/flox*^ embryos ^26^, the mutant lung buds still expressed Tbx4 in mesoderm (Supplementary Fig. S2a). Disruption of Wnt signaling in the mesoderm altered Tbx4 expression in the tracheal but not lung mesoderm, suggesting that Wnt-mediated mesodermal Tbx4 induction is a unique system in trachea development but not lung development.

We further found that the *Dermo1*^*Cre*^, *Ctnnb1*^*flox/flox*^ mutant exhibits tracheal cartilage agenesis. In the mutants, a periodic cartilage ring structure labeled with Sox9 failed to develop at E16.5, and circumferential SM bundles labeled with smooth muscle actin (SMA) were also malformed (Figs. 2e and e’) Therefore, mesodermal Wnt signaling is crucial for trachea mesenchymal development, particularly for tracheal cartilage development.

To determine whether Tbx4 is a direct or indirect target of canonical Wnt signaling in respiratory mesoderm, we explored Tcf/Lef binding sequences in the Tbx4 lung mesenchyme element (Tbx4-LME)^27, 28^. We identified 5 repeats of the Tcf/Lef binding sequence by using University of California Santa Cruz (UCSC) Genome browser and JASPAR (Fig. 2f)^29, 30^. These Tcf/Lef binding sequences were well conserved in all vertebrates except for fishes. These sequences are active cis-regulatory regions for H3K27Ac, H3K4me1 and p300 as determined by ChIP-seq and by chromatin accessibility.

Next, we sought to identify a source of Wnt ligands that initiates mesodermal Tbx4 expression. Due to the essential role of Wnt2 at early tracheal/lung development^4^, we conducted *in situ* hybridization for *Wnt2* and revealed transient expression of *Wnt2* in the ventrolateral mesoderm of the anterior foregut at E9.5, which was obviously reduced by E10.5 when Tbx4 was expressed (Figs. 2b and 3a). Wnt2 is most likely not involved in Tbx4 expression after E10.5. This observation prompted us to hypothesize that an endodermal-to-mesodermal interaction but not mesodermal autonomous induction is required for Tbx4 expression. To test this hypothesis, we generated *Shh*^*Cre*^, *Wls*^*flox/flox*^ mice, in which endodermal Wnt ligand secretion is inhibited by targeting *Wntless* (*Wls*) gene, which is essential for exocytosis of Wnt ligands^31^. This endoderm-specific deletion of *Wls* resulted in loss of Tbx4 expression in the mesoderm, but retained Nkx2.1 expression in the endoderm and *Wnt2* in the mesoderm (Figs. 3b and c)^31^, making these mice a phenocopy to *Dermo1*^*Cre*^, *Ctnnb1*^*flox/flox*^ mice (Fig. 2d). *Shh*^*Cre*^, *Wls*^*flox/flox*^ embryos also formed lung buds and expressed Tbx4 in the distal lung mesoderm (Supplementary Fig. S2b), supporting our idea that Wnt signaling in mesoderm mainly contributes to initiation of mesodermal development of the trachea, but not of the lung. These findings indicate that the endodermal Wnt ligands are sufficient for trachea mesodermal development. From these observations, we conclude that mesodermal Wnt2 activates endodermal canonical Wnt signaling which activates endodermal Wnt ligand expression independent of Nkx2.1. These Wnt ligands then induce endodermal-to-mesodermal canonical Wnt signaling to initiate mesodermal Tbx4 expression (Fig. 3d). These results also suggest that specification in the trachea endodermal lineage is not necessary for the initial induction of the tracheal mesoderm.

**Figure 3.**
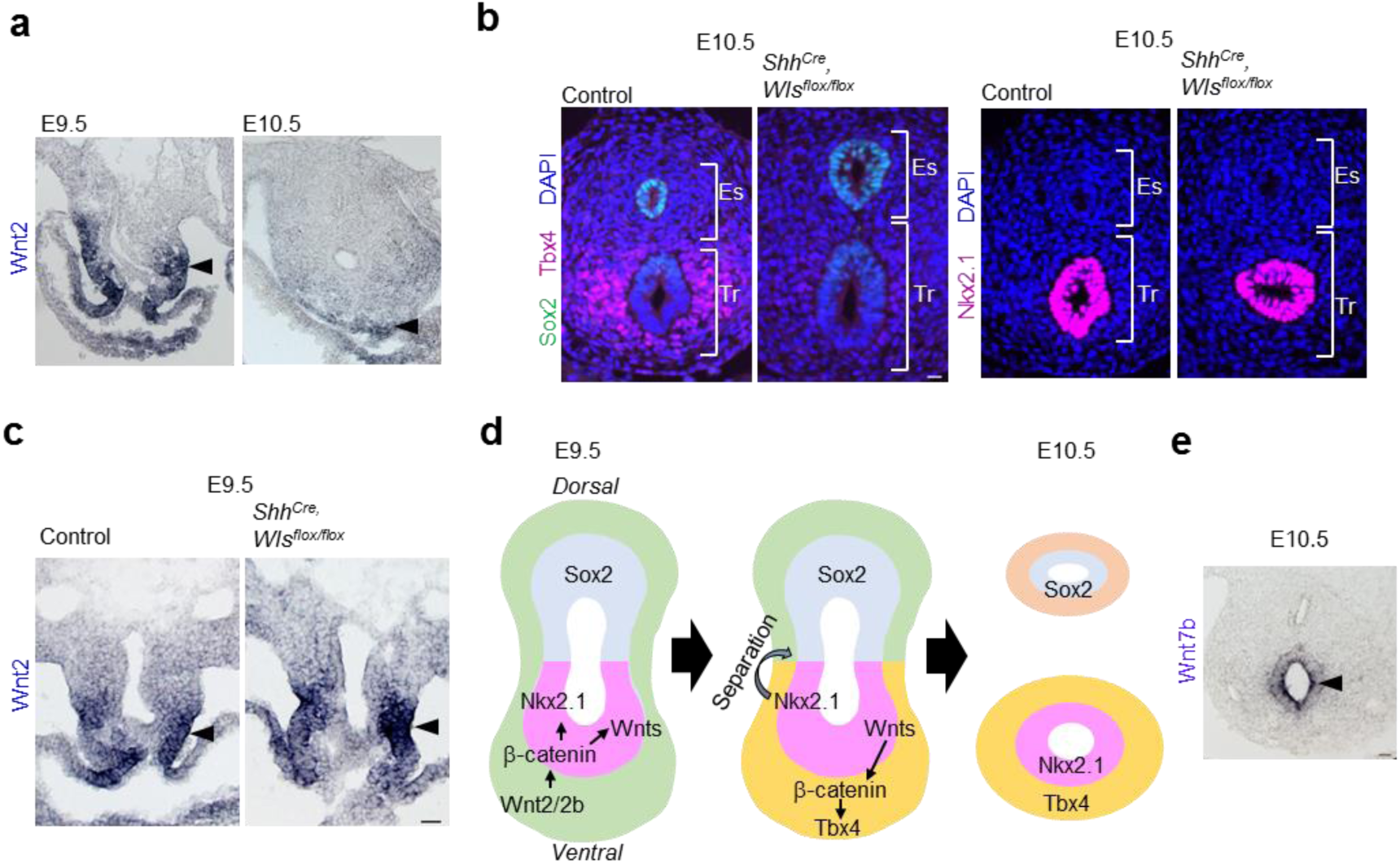
Endodermal Wnt ligands induce Tbx4 expression for tracheal mesoderm development of mouse trachea. **a**, *In situ* hybridization for *Wnt2* mRNA during tracheoesophageal segregation. Arrowheads indicate *Wnt2* expression in the ventrolateral mesoderm at E9.5 and E10.5. **b**, Transverse sections of *Shh*^*Cre*^, *Wls*^*flox/flox*^ mouse embryos and littermate controls at E10.5. Left panels show sections stained with Sox2 (*green*), Tbx4 (*magenta*), and DAPI (*blue*). Right panels show sections stained for Nkx2.1 (*magenta*) and DAPI (*blue*). **c**, *In situ* hybridization for Wnt2 mRNA in *Shh*^*Cre*^, *Wls*^*flox/flox*^ mouse embryos and littermate controls at E9.5. Arrowheads indicate Wnt2 expression in the ventrolateral mesoderm. **d**, Refined model of tracheoesophageal segregation and tracheal mesodermal differentiation **e**, *In situ* hybridization for Wnt7b mRNA in mouse embryo at E10.5. Arrowhead indicates Wnt7b^+^ cells. Eso; Esophagus, Lu; Lung, Tr; Trachea. Scale bar; 40 μm (a-c), 50 μm (e).

In the developing mouse trachea, several Wnt ligands are expressed in the endoderm between E11.5 to E13.5, such as Wnt3a, 4, 5a, 6, 7b, 11 and 16^26, 31^. Current single cell RNA-seq data have shown the presence of several Wnt ligands including Wnt4, 5a, 5b, 6 and 7b in the respiratory endoderm of mouse embryos at E9.5 (Han et al., back-to-back). We performed *in situ* hybridization (ISH) against these Wnt ligands at E10.5 to determine the particular ligand inducing Tbx4 expression in trachea development (Fig. 3e, Supplementary Fig. S4). Wnt4 was expressed in esophageal mesoderm and barely detected in tracheal endoderm. Wnt5a, 5b and 6 were detected in both the endoderm and mesoderm of the trachea. More importantly, Wnt7b was abundantly expressed in tracheal endoderm, suggesting that Wnt7b might be responsible for the ensuing induction of mesodermal Tbx4 expression

To examine whether Wnt signaling is capable of initiating the differentiation of naïve mesodermal cells to Tbx4^+^ trachea mesodermal cells in *vitro*, we established a protocol for lateral plate mesoderm (LPM) induction from mouse ESCs by refining the published protocol for LPM induction from human pluripotent stem cells^32^. Because mouse and human ESCs show different states called naïve and primed, which correspond to pre- and post-implantation epiblasts, respectively, we converted mouse ESCs (mESCs) into an epiblast ‘primed’ state that led to middle-primitive streak (mid-PS) cells^33^. These mid-PS cells were then differentiated into LPM cells (Fig. 4a). At day 5, LPM induction was confirmed by immunocytochemistry (ICC) for Foxf1 and Gata4 which are known to be expressed in LPM including splanchnic mesoderm^34, 35^ (Fig. 4b), and showed that 89% of total cells were Foxf1^+^ LPM. qRT-PCR also showed obvious upregulation of LPM marker genes such as Foxf1, Gata4, Hoxb6, Prrx1, and Bmp4 (Figs. 4c and e). Given that previous mouse genetic studies have identified Bmp4 as a crucial regulator of trachea development^6, 36^, we tested whether canonical Wnt and Bmp4 signaling are sufficient to direct the differentiation of LPM into the tracheal mesoderm (Foxf1^+^/Tbx4^+^). mESC-derived LPM cells were cultured with CHIR99021, a GSK3β inhibitor to stabilize β-catenin and activate canonical Wnt signaling, and Bmp4. At day 6, 89% of total cells became double positive for Foxf1^+^ and Tbx4^+^. qRT-PCR further demonstrated elevated expression of tracheal marker genes such as Tbx5, Wnt2, Bmp4 in addition to Tbx4 (Figs. 4d and e). To further confirm the respiratory characteristics of these cells, we took advantage of the 5 repeats of Tcf/lef binding sequences in Tbx4-LME, which we described in Fig. 2f. We established a luciferase reporter assay by reporter plasmids that express luciferase under the control of Tbx4-LME (Fig. 4f). The reporter plasmid was transfected into mESC-derived LPM and luciferase activity was assessed during differentiation. After 24 hours (at day 6), the luciferase activity increased in the presence of CHIR99021 (Fig. 4g). Importantly, the mutated reporter, in which all Tcf/Lef binding sequences were changed to random sequences (Figs. 2f and 4f), did not respond to CHIR99021. These results determined that the mESC-derived cells were differentiated into proper tracheal mesoderm at day 6.

**Figure 4.**
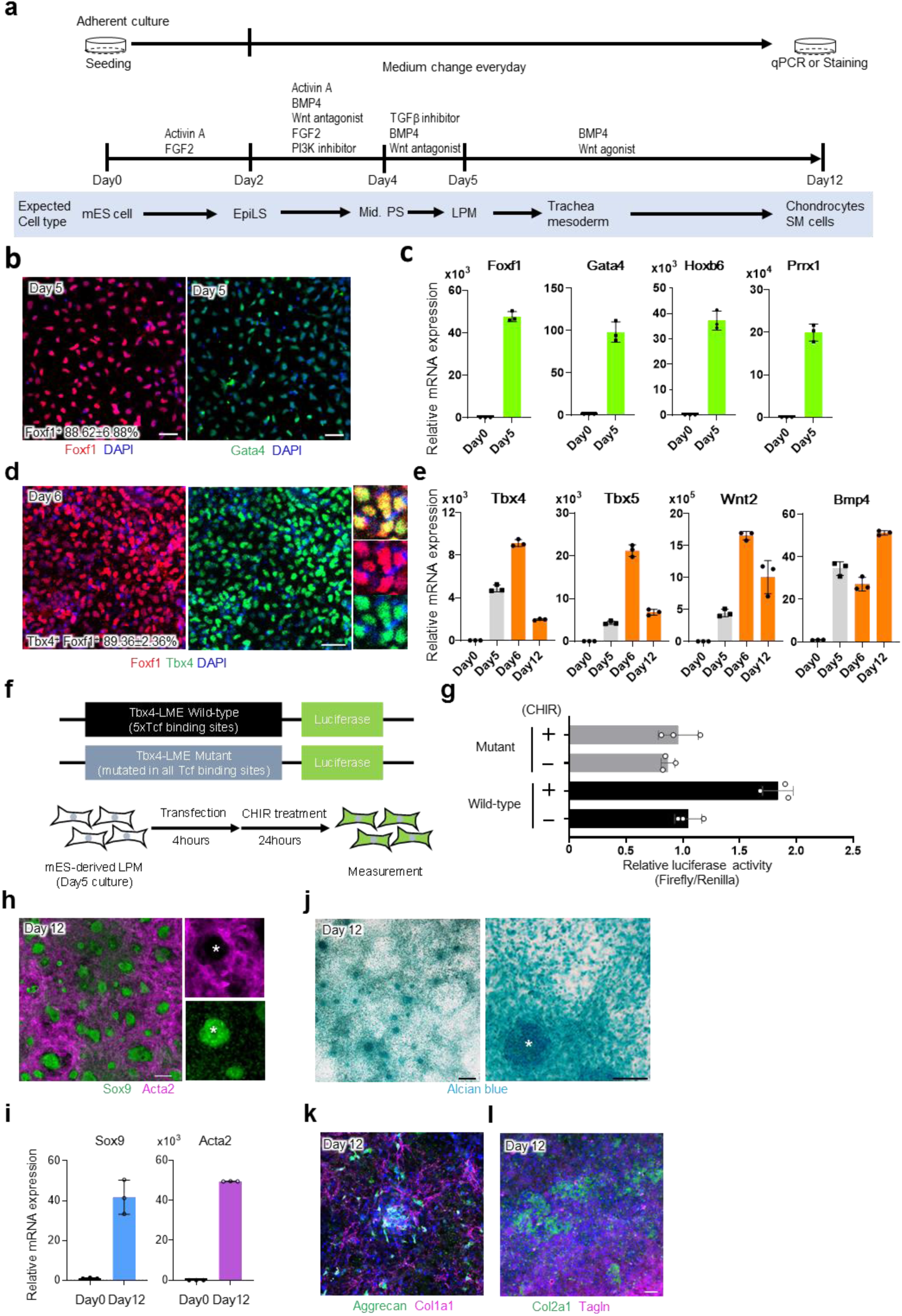
Generation of trachea mesodermal cells and chondrocytes from mouse ESCs *in vitro.* **a**, Experimental design to generate tracheal mesoderm from mESCs. **b**, Differentiating cells from mESCs at day 5. Cells were stained for Foxf1 (*red*) and Gata4 (*green*), respectively. **c**, Results of qRT-PCR for LPM marker expression of mESC-derived LPM at day 0 and 5. **d**, Differentiating cells from mESCs at day 6. Cells were stained for Tbx4 (*green*) and Foxf1 (*red*). **e**, Results of qRT-PCR for respiratory mesoderm marker expression of mESC-derived tracheal mesodermal cells during differentiation. **f**, Diagram showing the constructs utilized in luciferase experiments containing *Tbx4* LME wild-type containing five Tcf/Lef-binding sites (5xTcf) and Tbx4-LME Mutant. mESC-derived LPM cells transfected with wild type or mutant *Tbx4* LME during 4 hrs following by respiratory induction in presence or absence of CHIR99021. **g**, Luciferase assay examining the activation of *Tbx4* LME wt and mutant in response to 3μM CHIR99021 **h** Differentiating cells from mESCs at day 12. Cells were stained for Acta2 (*magenta*) and Sox9 (*green*). The asterisk indicates Sox9^+^/SMA^-^ chondrocyte aggregates. **i**, Results of qRT-PCR for *Sox9* and *Acta2* expression of hESC-derived tracheal mesodermal cells at day0 and 12. **j**, Differentiating cells from mESCs at day 12. Chondrocytes were stained with Alcian blue. The asterisk indicates one of the chondrocyte aggregates. **k**, Differentiating cells from mESCs at day 12. Cells were stained for Col1a1 (*magenta*) and Aggrecan (*green*). **l**, Differentiating cells from mESCs at day 12. Cells were stained for Tagln (*magenta*) and Col2a1 (*green*). Each column shows the mean with S.D. (n=3). Scale bar; 50μm. Source data for b, c, d, e, g, i are provided in Source data file.

Because tracheal mesenchyme includes cartilage and smooth muscle, we wondered whether our protocol induces mESC to differentiate into these tissues. At day 12, Sox9^+^ aggregated cell masses positive for Alcian blue staining appeared on the dish, indicative of chondrocytes (Figs. 4h-j). Smooth muscle cells (SMA^+^ cells) concurrently appeared to show fibroblastic morphology and filled the spaces not filled by the Sox9^+^ cells (Figs. 4h and i). Other chondrogenic markers (Aggrecan, Collagen2a1, Sox5/6, Epiphycan) and smooth muscle markers (Tagln, Collagen1a1) were also present in the differentiated cells (Figs. 4k, l and Supplementary Fig. S6a). These data suggest that the mESC-derived tracheal mesoderm is able to develop into tracheal mesenchyme, including chondrocytes and smooth muscle cells.

Finally, we tested the role of Wnt signaling in the human tracheal mesoderm using human ESCs (hESCs). Human LPM induction was performed by following an established protocol^32^ (Fig. 5a). Subsequently, the cells were directed to tracheal mesoderm by using CHIR99021 and BMP4. For validating hESC-derived LPM, we checked the common LPM markers at day 2 and confirmed that these markers were abundantly expressed in the LPM (Figs. 5b, c and Supplementary Fig. S7j). Immunostaining determined that 95% of the total cells expressed FOXF1 at day 2 (Fig. 5b). Because Tbx4 is also expressed in the limbs and other fetal mouse tissues^28^, we sought additional genetic markers for the tracheal mesoderm. We searched the single-cell transcriptomics dataset of the developing splanchnic mesoderm at E9.5 and identified *Nkx6.1* as a marker for mesodermal cells surrounding the trachea, lung and esophagus (Han et al., co-submitted). We performed immunostaining and found that Nkx6.1 was expressed in tracheal and esophageal mesenchyme throughout development (Supplementary Fig. S5). Of note, Nkx6.1 was expressed in esophageal and dorsal tracheal mesenchyme but not ventral trachea, which enabled us to define three subtypes of tracheal-esophageal mesenchyme based on the combination of Tbx4 and Nkx6.1 expression (i.e. Tbx4^+^/Nkx6.1^+^; dorsal tracheal mesenchyme, Tbx4^+^/Nkx6.1^-^; ventral tracheal mesenchyme, Tbx4^-^ /Nkx6.1^+^; esophageal mesenchyme) (Supplementary Fig. S5). Having characterized the subtypes of tracheal-esophageal mesoderm *in vivo*, the expression of *TBX4* and *NKX6.1* in the hESC-derived tracheal mesoderm was examined by ICC and qRT-PCR. Although *TBX4* was induced in a Wnt agonist dose-dependent manner, *NKX6.1* expression was not significantly elevated (Supplementary Figs. S7a, b), suggesting that human trachea mesodermal development requires an additional factor to become more *in vivo*-like. Because the ventral LPM is exposed to SHH in addition to Wnt and Bmp4 during tracheoesophageal segregation^37, 38^, we assessed whether the SHH agonist (PMA; purmorphamine) can improve differentiation from hESC-derived LPM cells into the tracheal mesoderm. As expected, both *TBX4* and *NKX6.1* expression was upregulated by the SHH agonist (Supplementary Figs. S7c, d). After day 5, the differentiating cells also expressed respiratory markers such as TBX4, TBX5, WNT2, BMP4 and NKX6.1 (Figs. 5d-h, Supplementary Fig. S7j). In this culture condition, CHIR99021 enhanced the expression of *TBX4* and *NKX6.1* genes in a dose-dependent manner (Supplementary Figs. S7e, f). We further estimated the efficiency of the induction by immunostaining for TBX4 and FOXF1, and then confirmed that 83% of total cells were TBX4^+^/FOXF1^+^ double positive cells at day 5 (Fig. 5d). At day 10, NKX6.1 expression was clearly elevated, and 30.3% of the total cells became TBX4^+^/NKX6.1^+^ double positive (Figs. 5e-g) while 18.0% were TBX4^+^/NKX6.1^-^ (Fig. 5g). These data suggest that the half of the cells induced with our protocol are trachea mesodermal cells. Further extended culture induced SOX9^+^ aggregates which were positive for Alcian blue staining in a Wnt activity-dependent manner (Figs. 5h-l). Likewise in mESC-derived cells, ACTA2^+^ smooth muscle-like fibroblastic cells occupied Sox9^-^ region (Fig. 5i). These cells also expressed chondrogenic markers and smooth muscle cell markers (Figs. 5k, l and Supplementary Fig. S6b).

**Figure 5.**
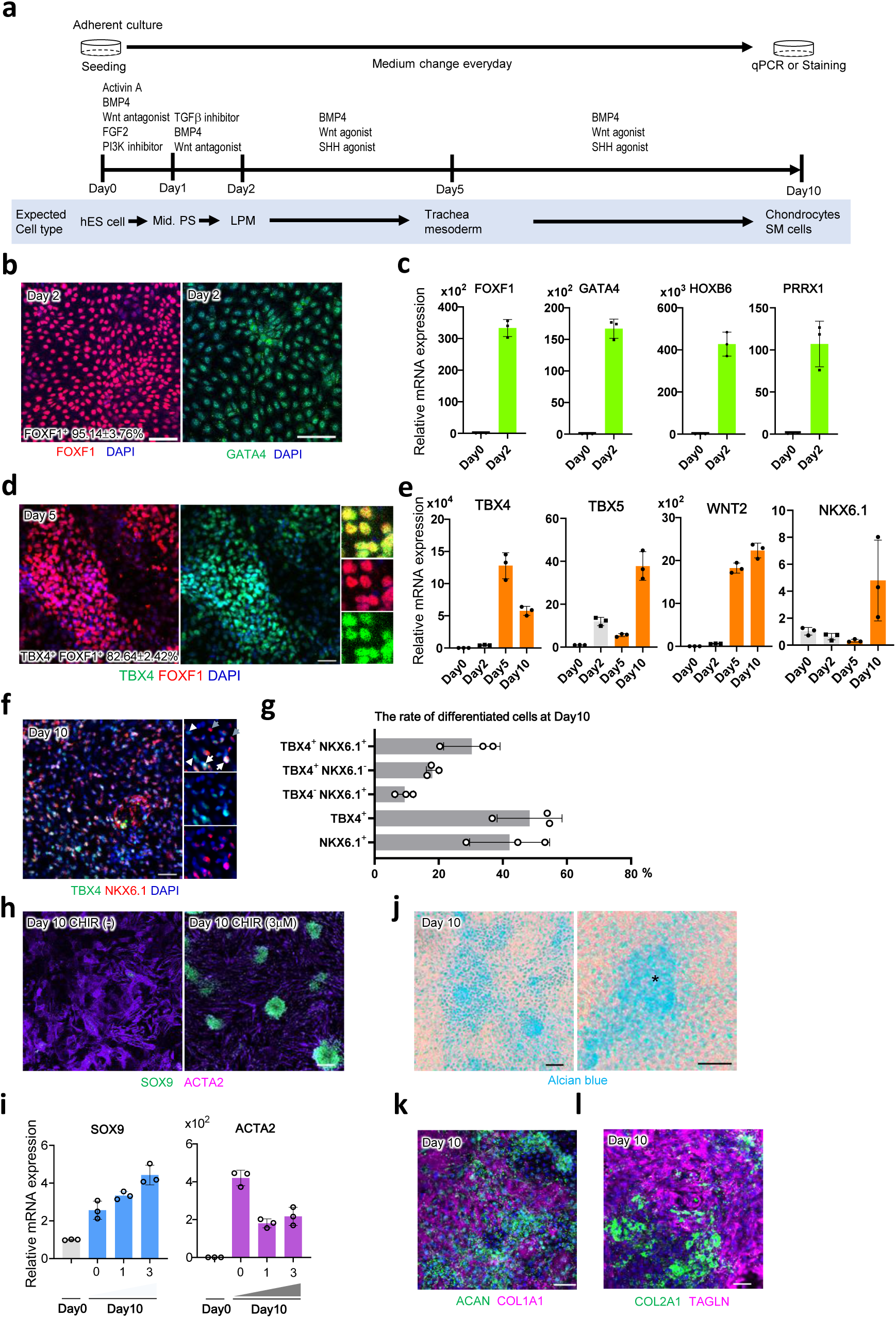
Generation of trachea mesodermal cells and chondrocytes from human ESCs *in vitro.* **a**, Experimental design to generate tracheal mesoderm from hESCs **b**, Differentiating cells from hESCs at day 2. Cells were stained for FOXF1 (*red*) and GATA4 (*green*), respectively. **c**, Results of qRT-PCR for LPM marker expression of hESC-derived LPM at day2 **d**, Differentiating cells from hESCs at day5. Cells were stained for TBX4 (*green*) and FOXF1 (*red*). **e**, Results of qRT-PCR for respiratory mesoderm marker expression of hESC-derived tracheal mesodermal cells during differentiation. **f**, Differentiating cells from hESCs at day10. Cells were stained for TBX4 (*green*) and NKX6.1 (*red*). White arrows indicate TBX4^+^/NKX6.1^+^ mesodermal cells. White arrowheads indicate TBX4^+^/NKX6.1^-^ mesodermal cells. Grey arrows indicate TBX4^-^ /NKX6.1^+^ mesodermal cells. **g**, The rate of differentiated cells at day10. **h**, Differentiating cells from hESCs with or without CHIR99021 treatment at day 10. Cells were stained for ACTA2 (*magenta*) and SOX9 (*green*). **i**, Results of qRT-PCR for *SOX9* and *ACTA2* expression of hESC-derived trachea mesodermal cells at day 10 with different doses of CHIR99021. **j**, Differentiating cells from hESCs at day 10. Chondrocytes were stained with Alcian blue. The asterisk indicates one of the chondrocyte aggregates. **k**, Differentiating cells from hESCs at day 10. Cells were stained for COL1A1 (*magenta*) and AGGRECAN (*green*). **l**, Differentiating cells from hESCs at day 10. Cells were stained for TAGLN (*magenta*) and COL2A1 (*green*). Each column shows the mean with S.D. (n=3). Scale bar; 50μm (d, f, h, j, k, l), 100μm (b) Source data for b, c, d, e, g, i are provided in Source data file.

In this culture system, the removal of BMP4 from the growth factor cocktail did not affect differentiation, implying that exogenous BMP4 activation is dispensable (Supplementary Fig. S7g-i). Because of the obvious upregulation of the endogenous *BMP4* gene in the hESC-derived LPM by day 2, endogenous BMP4 may be enough to induce tracheal mesoderm and chondrocytes (Supplementary Fig. S7j). Taken together, these data suggest that Wnt signaling plays a unique role in driving differentiation into tracheal mesoderm and chondrocytes from the LPM, which is conserved between mice and human.

This study demonstrated that endodermal-to-mesodermal canonical Wnt signaling is the cue that initiates trachea mesodermal development in developing mouse embryos, which is independent of the previously known Nkx2.1-mediated respiratory tissue development. Based on our knowledge of developmental biology, we successfully generated tracheal mesoderm and chondrocytes from mouse and human ESCs. In our protocol, we stimulated ESC-derived LPM with Wnt, Bmp and SHH signaling to mimic spatial information of the ventral anterior foregut. For induction of respiratory endoderm, Wnt, Bmp and Fgf signaling are required to direct cells in anterior foregut to differentiate into the respiratory lineage^9-11^. Thus, Wnt and Bmp signaling are conserved factors that provide spatial information, while Fgf and SHH are required in endoderm and mesoderm induction, respectively, reflecting the unique signaling pathways in each tissue. Mesoderm induction may need fewer exogenous growth factors because the mesodermal cells themselves are a source of spatial information, such as BMP4 in our protocol.

In this study, we were unable to perform tissue-specific targeting for trachea endodermal or mesodermal cells because of multiple Cre-expression patterns in Shh-Cre and Dermo1-Cre mouse lines. For example, Shh is also expressed in the notochord and ventral neural tube^39^. Future studies with analyses of respiratory tissue-specific Cre lines would strengthen the evidence demonstrating that mutual interaction between respiratory endoderm and mesoderm is required for the induction of trachea development.

*Dermo1*^*Cre*^, *Ctnnb1*^*flox/flox*^ mutants display a tracheal cartilage agenesis phenotype. Due to the multiple functions of Ctnnb1 in transcriptional regulation and cellular adhesion, however, it is possible that Ctnnb1 knockout affects not only Wnt-mediated transcriptional regulation but also mesenchymal cell-cell adhesion^24^. To exclude this possibility, we examined the distribution of CDH2 as an adhesive molecule in tracheal mesoderm. CDH2 expressions in the ventral half of tracheal mesoderm were indistinguishable between control and *Dermo1*^*Cre*^, *Ctnnb1*^*flox/flox*^ embryos (Supplementary Fig. S3). Furthermore, our luciferase assay revealed that respiratory mesenchyme specific cis-regulatory region of Tbx4 is stimulated by CHIR99021 through Tcf/Lef binding elements in the developing tracheal mesoderm. These findings suggest that Wnt signaling-mediated transcriptional regulation is important for the induction of tracheal mesoderm.

Recently, Han et al. delineated mesodermal development during organ bud specification using single-cell transcriptomics analyses of mouse embryos from E8.5 to E9.5 (Han et al., co-submitted to *Nature communications*). Based on the trajectory of cell fates and signal activation, this group also generated organ-specific mesoderm, including respiratory mesoderm, from hESCs, thereby determining that Wnt, BMP4, SHH and retinoic acid direct differentiation of hESC-derived splanchnic mesoderm to respiratory mesoderm, supporting our current findings.

These culture methods could be a strong tool to study human organogenesis and the aetiology of TEA and TA, as well as to provide cellular resource for human tracheal tissue repair.

## Methods

### Mice

All mouse experiments were approved by the Institutional Animal Care and Use Committee of RIKEN Kobe Branch. Mice were handled in accordance with the ethics guidelines of the institute. *Nkx2.1*^*null*^, *Shh*^*Cre*^, *Dermo1*^*Cre*^, *Ctnnb1*^*flox/flox*^, *Wls*^*flox/flox*^ mice were previously generated^18, 23, 40-42^.

In all experiments, at least 3 embryos from more than 2 littermates were analyzed. All attempts for replicate were successful. Sample size was not estimated by statistical methods. No data was excluded in this study. All control and mutant embryos were analyzed. No blinding was done in this study.

### Immunostaining

Mouse embryos were fixed by 4% Paraformaldehyde/PBS (PFA) at 4°C overnight. Specimens were dehydrated by ethanol gradient and embedded in paraffin. Paraffin sections (6-μm) were de-paraffinized and rehydrated for staining. Detailed procedure and antibodies of each staining were listed in Supplementary Table 1.

### In situ hybridization

Mouse embryos were fixed with 4%PFA/PBS at 4°C overnight, and then tracheas were dissected. Specimens were incubated in sucrose gradient (10, 20, 30%) and embedded in OCT compound. Frozen sections (12-μm) were subjected to *in situ* hybridization. For Wnt2, 4, 5a, 7b probe construction, cDNA fragments were amplified by primers listed in Supplementary Table 2. These cDNA fragments were subcloned into pBluscript SK+ at *Eco*RI and *Sal*I sites. For Wnt5b and 6 probes, pSPROT1-Wnt5b (MCH085322) and pSPROT1-Wnt6 (MCH000524) were linearized at *Sal*I sites, The NIA/NIH Mouse 15K and 7.4K cDNA Clones were provided by the RIKEN BRC^43-45^. Antisense cRNA transcripts were synthesized with DIG labeling mix (Roche Life Science) and T3 or SP6 RNA polymerase (New England Biolabs Inc.). Slides were permeabilized in 0.1% Triton-X100/PBS for 30min and blocked in acetylation buffer. After pre-hybridization, slides were hybridized with 500ng/ml of DIG-labeled cRNA probes overnight at 65°C. After washing with SSC, slides were incubated with anti-DIG-AP antibodies (1:1000, Roche Life Science, 11093274910). Sections were colored with BM-purple (Roche Life Science, 11442074001).

For RNAscope experiments, the RNAscope Multiplex Fluorescent v2 assays (Advanced Cell Diagnostics, 323110) were used. The detailed procedure and probes were listed on Supplementary Table 3

### Cell culture

For mesodermal differentiation from mES cells, C57BL/6J-Chr 12A/J/NaJ AC464/GrsJ mES cells (The Jackson Laboratory) and EB3 cells (AES0139, RIKEN BioResorce Center) were used. C57BL/6J-Chr 12A/J/NaJ AC464/GrsJ mES cells were kindly provided by Kentaro Iwasawa and Takanori Takebe (Center for Stem Cell & Organoid Medicine (CuSTOM), Perinatal Institute, Division of Gastroenterology, Hepatology and Nutrition, Cincinnati Children’s Hospital, Cincinnati). EB3 was kindly provided by Dr. Hitoshi Niwa (Department of Pluripotent Stem Cell Biology, Institute of Molecular Embryology and Genetics in Kumamoto University)^46, 47^. Cells were maintained in 2i + leukemia inhibitory factor (LIF) media (1,000 units/ml LIF, 0.4μM PD0325901, 3μM CHIR99021 in N2B27 medium) on ornithine-laminin coated-dishes^33^. For mesodermal differentiation of mouse ES cells, cells were digested by TrypLE express (Thermo Fisher Scientific, 12604013) and seeded onto Matrigel-coated 12 well plate. EpiLC were induced by EpiLC differentiation medium (1% knockout serum, 20ng/ml Activin A, 12ng/ml FGF2, and 10μM Y27632 in N2B27 Medium)^33^ for 2 days. Lateral plate mesoderm was established by Loh’s protocol with some modification^32^. EpiLC cells were digested by TrypLE express to single cells and seeded onto Matrigel-coated 12 well plate at the density of 6×10^5^ cells/well. The cells around middle primitive streak was induced by LPM D2 medium composed of 2% B27 Supplement Serum free (Thermo Fisher Scientific, 17504044), 1 x GlutaMax (Thermo Fisher Scientific, 35050061), 20ng/ml basic FGF (Peprotech, AF-100-18B), 6μM CHIR99021 (Sigma Aldrich, SML1046), 40ng/ml BMP4 (R&D Systems, 5020-BP-010), 10ng/ml Activin A (Peprotech, PEP-120-14-10), 10μM Y27632 (Sigma Aldrich, Y0503) in Advanced DMEM (Thermo Fisher Scientific, 12491015) for 48 hours. After that, LPM was induced by LPM D4 medium composed of 2% B27 Supplement Serum free, 1 x GlutaMax, 2μM XAV939 (Sigma Aldrich, X3004), 2μM SB431542 (Merck, 616461), 30ng/ml human recombinant BMP4 in Advanced DMEM for 24 hours. At Day 5, respiratory mesenchyme was induced by Day5 medium composed of 2% B27 Supplement Serum free, 1 x GlutaMax, 1μM CHIR99021, 10ng/ml BMP4. Medium were freshly renewed every day.

H1 (NIHhESC-10-0043 and NIHhESC-10-0062), human embryonic stem cell, was provided by Cincinnati children’s hospital medical center Pluripotent Stem Cell Facility. Cells were maintained in mTeSR1 medium (Stem Cell Technologies) on Matrigel-coated plate. For differentiation of H1 cells to mesodermal cells, confluent cells were digested by Accutase to single cells and seeded onto Geltrex-coated 12well plate at the dilution of 1:20 – 1:18 in mTeSR1 with 1uM Thiazovivin (Tocris). Next day, the cells around middle primitive streak were induced by cocktails of 6μM CHIR99021 (Sigma Aldrich, SML1046), 40ng/ml BMP4 (R&D Systems, 5020-BP-010), 30ng/ml Activin A (Cell Guidance Systems), 20ng/ml basic FGF (Thermo Fisher Scientific) and 100nM PIK90 (EMD Millipore) in Advanced DMEM/F12 including 2% B27 Supplement minus vitamin A, 1% N2 Supplement, 10uM Hepes, 100UI/mL Penicillin/Streptomycin, 2mM L-glutamine for 24 hours. After that, LPM was induced by LPM D2 medium composed of 1μM Wnt C59 (Cellagen Technologies), 1μM A83-01 (Tocris), 30ng/ml human recombinant BMP4 in Advanced DMEM/F12 including 2% B27 Supplement minus V. A., 1 x N2 Supplement, 10uM Hepes, 100UI/mL Penicillin/Streptomycin, 2mM L-glutamine for 24 hours. To generate respiratory mesenchyme, we combined 3uM CHIR99021, 2uM Purmorphamine (Tocris), and 10ng/ml Bmp4 in Advanced DMEM/F12 medium including 2% B27 Supplement Serum free, 1 x N2 Supplement, 10uM Hepes, 100UI/mL Penicillin/Streptomycin, 2mM L-glutamine from Day 2 to Day10. Medium was freshly renewed everyday

### Immunocytochemistry

At differentiating process, cells were fixed by 4% PFA for 10 minutes at room temperature. For intracellular staining, cells were permeabilized by 0.2% TritonX-100/PBS for 10 minutes at room temperature. After blocking the cells with 5% normal donkey serum, cells were incubated with primary antibodies overnight at 4°C. Then, cells were incubated with secondary antibodies for 1hr at room temperature. Detailed procedure and antibodies of each staining were listed in Supplementary Table 4.

### Luciferase reporter assay

The fraction of mouse *Tbx4*-lung mesenchyme specific enhancer (LME) (mm10, chr11:85,893,703-85,894,206, GenScript, ID U3154EL200-3)^27^ or *Tbx4*-LME containing putative Tcf/Lef sites mutated (GenScript, ID U3154EL200-6) were synthesized and cloned into pGL4.23 (luc2/minP) vector (promega).

mESC-derived LPM cells were transfected at day 5 in 150µl of Opti-MEM (Thermo Fisher Scientific, 31985088) with 2µl of Lipofectamine Stem (Thermo Fisher Scientific, STEM00003) and 1µg of pGL4.23 (luc2/minP) containing a fraction of mouse *Tbx4*-LME or *Tbx4*-LME containing mutated Tcf/Lef sites. Four hours after transfection the tracheal mesenchyme was induced using day5 medium in presence or absence of Wnt agonist (3μM CHIR99021) and cells were cultured for 24 hours and then lysed and assayed using Dual-Luciferase Reporter Assay System (Promega, E1980).

### Alcian blue staining

Cells were fixed in 4% PFA/PBS for 10 minute at room temperature. After washing with PBS, cells were incubated with 3% acetic acid for 3 minutes and then stained with 1% alcian blue/3% acetic acid for 20 minutes.

### Quantitative RT-PCR

Total mRNA was isolated by using the Nucleospin kit (TaKaRa, 740955) according to manufacturer’s instruction. cDNA was synthesized by Super™Script™ VILO cDNA synthesis kit (Thermo Fisher Scientific, 11754050). qPCR was performed by PowerUp™ SYBR™ Green Master Mix on QuantStudio 3 or 6. Primer sequences were listed on Supplementary Table 5 and 6. Data are expressed as a Fold Change and were normalized with undifferentiated cells expression.

### Statistical analyses

Statistical analyses were performed with Excel2013 (Microsoft) or PRISM8 (GraphPad software). For multiple comparison, one-way ANOVA and Tukey’s methods were applied. For paired comparison, statistic significance was determined by F-test and Student’s or Welch’s two-tailed t test.

## Acknowledgements

We thank Hinako M Takase and Hiroshi Hamada for Wntless conditional flox mice, Masatoshi Takeichi for Ctnnb1 conditional flox mice, and Animal Resource Development Unit. Hiroshi Niwa kindly provided EB3, mouse ES cells. Kentaro Iwasawa and Takanori Takebe kindly provided C57BL/6 mouse ES cell line. We are grateful to Debora Sinner for the help on RNAscope experiment. We thank Scott Rankin for the help on the enhancer analyses of Tbx4 genes. We also thank Shunsuke Mori and Mototsugu Eiraku for Tbx4 antibody. We thank Yuka Noda and David Luedeke for general technical support. We also thank Shigeo Hayashi for primary reading.

These studies are supported by the funding from Grants-in-Aid for Scientific Research (B)(17H04185)(20H03693)(M.M.), Young Scientists (17K15133 and 19K16156)(K.K.), Promotion of Joint International Research (A) (18KK0423)(K.K.) of the Ministry of Education, Culture, Sports, Science and Technology, Japan, and from The Takeda Science Foundation for the Life Science (M.M.), and The Uehara Memorial Foundation (K.K.). Partially supported by grant NICHD P01HD093363 to AMZ.

## Author contributions

K.K., and M.M. designed the project and performed experiments with the aid of A.L.M., A.Y., C. M., and K.T.F. AMZ analysed single cell transcriptomics for definitive endoderm and splanchnic mesoderm. A.L.M. performed enhancer analyses of Tbx4 gene and supported human ES cell experiments, A.Y. supported mouse experiments. K.K., K.T. F. and C.M. performed mouse ES cell experiment. C. A. and M.H. contributed to mouse and human embryonic-stem cell-based lateral plate mesoderm induction and differentiation experiments.

K.K. and M.M. wrote the manuscript with the contribution of all authors.

## Data availability

The authors declare that all data supporting the findings of this study are available within the article and its Supplementary Information files or from the corresponding author upon reasonable request. The Source data underlying Figs. 4b-e, 4g, 4i, 5b-e, 5g, 5i and Supplementary Figs. S6 and S7 were provided as a Source data file.

The datasets generated during the current studies are available in the System Science of Biological Dynamics (SSBD) database (http://ssbd.qbic.riken.jp/).

