## Supplementary Info for "Bidirectional Wnt signaling between endoderm and mesoderm confer tracheal identity in mouse and human"

**by Kishimoto et al.**

Supplementary Figure S1

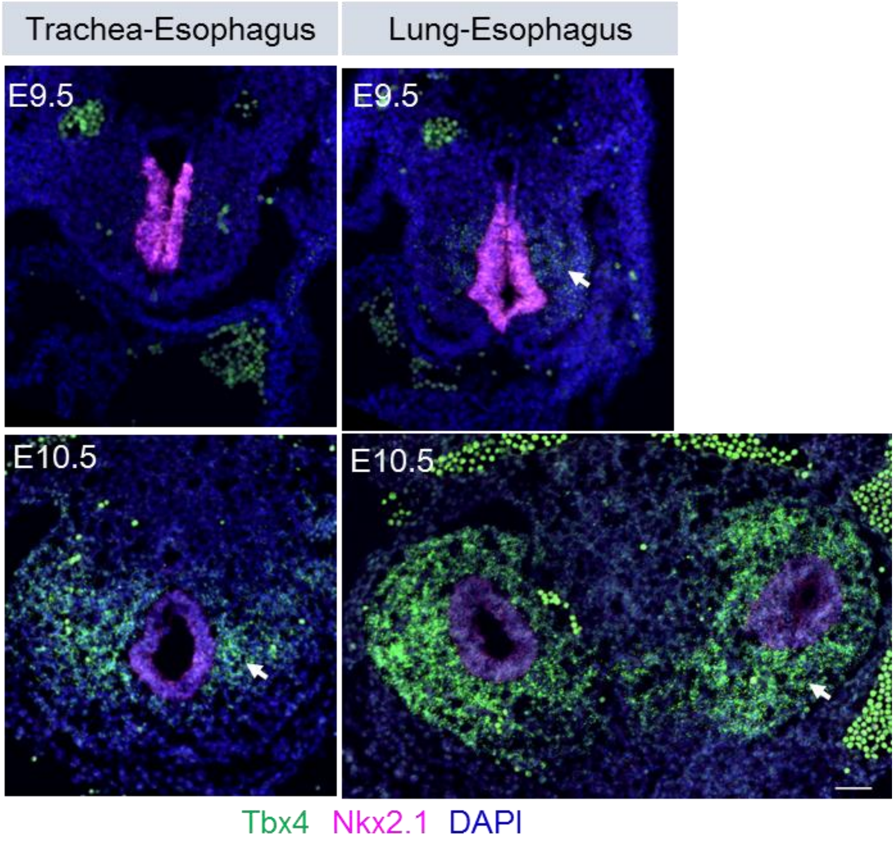


**Supplementary Figure S1 Tbx4 mRNA expression during tracheal-esophageal segregation at E9.5 and 10.5.**

RNAscope *in situ* hybridization for *Tbx4* mRNA during tracheoesophageal segregation. Sections were stained by Tbx4 (*green*), Nkx2.1 (*magenta*), and DAPI (*blue*). Arrows indicate Tbx4$^{+}$ cells. Scale bar; 50µm

Supplementary Figure S2

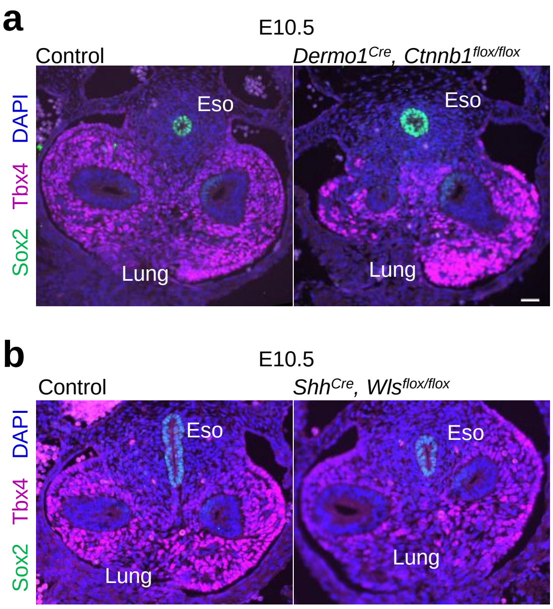


**Supplementary Figure S2 Dispensable role of mesodermal Wnt signaling in Tbx4 expression during lung development.**

**a**, Transverse sections of *Dermo1*$^{Cre}$, *Ctnnb1*$^{flox/flox}$ mouse embryos and littermate controls. Sections were stained for Sox2 (*green*), Tbx4 (*magenta*), and DAPI (*blue*).

**b**, Transverse sections of *Shh*$^{Cre}$, *Wls*$^{flox/flox}$ mouse embryos and littermate controls. Sections were stained for Sox2 (*green*), Tbx4 (*magenta*), and DAPI (*blue*).

Eso; Oesophagus, Tr; Trachea, Tr-E; Tracheoesophageal tubes

Scale bar; 40 μm.

Supplementary Figure S3

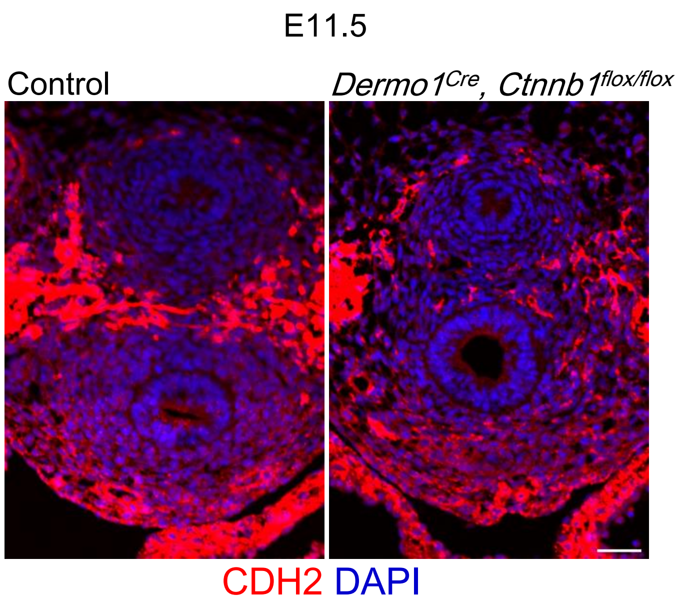


**Supplementary Figure S3 Distribution of CDH2 in trachea and esophagus in *Dermo1*$^{Cre}$, *Ctnnb1*$^{flox/flox}$ embryos at E11.5**

Transverse sections of *Dermo1*$^{Cre}$, *Ctnnb1*$^{flox/flox}$ mouse embryos and littermate controls. Sections were stained for CDH2 (*red*), and DAPI (*blue*). Scale bar; 40μm

Supplementary Figure S4

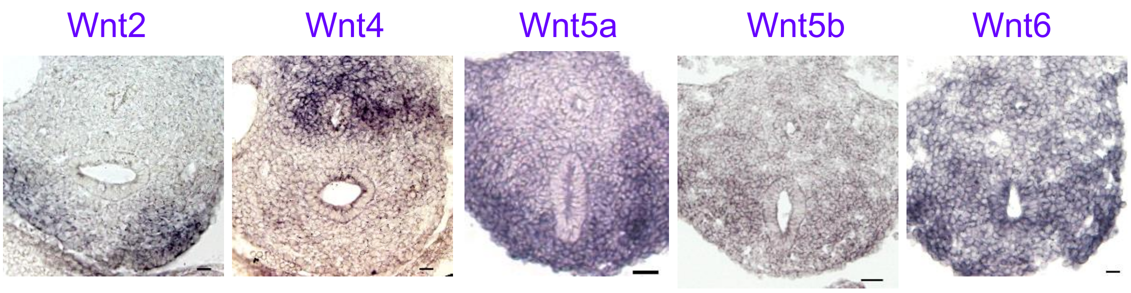


**Supplementary Figure S4 Wnt ligand expression in trachea and esophagus at E10.5**

*In situ* hybridization for Wnt2/4/5a/5b/6 mRNA in mouse embryos at E10.5.

Scale bar; 50µm

Supplementary Figure S5

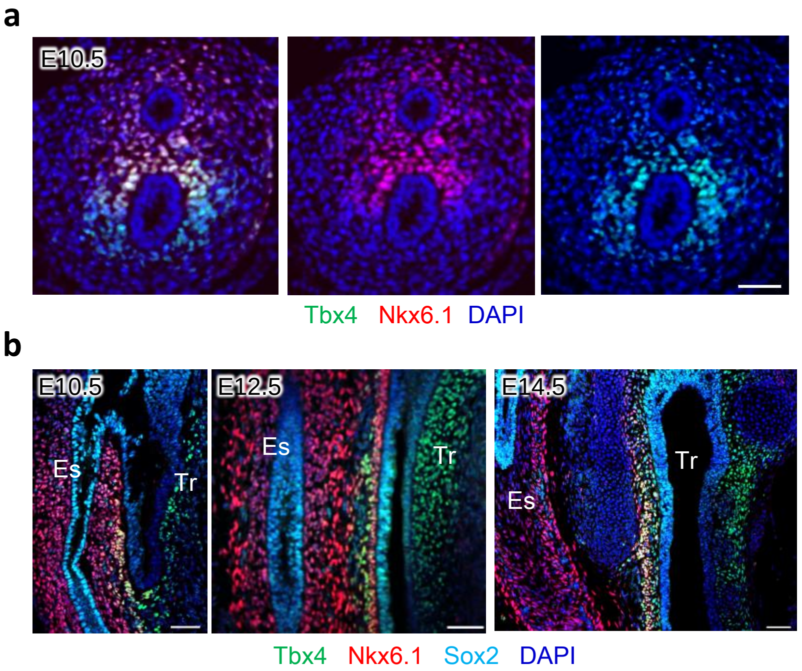


**Supplementary Figure S5 Nkx6.1 and Tbx4 expression during trachea development.**

**a**, Transverse sections of mouse embryo at E10.5. Sections were stained for Tbx4 (*green*), Nkx6.1 (*red*), and DAPI (*blue*).

**b**, Sagittal sections of mouse embryo from E10.5 to 14.5. Sections were stained for Tbx4 (*green*), Nkx6.1 (*red*), Sox2 (*cyan*) and DAPI (*blue*).

Es; Esophagus, Tr; Trachea, Scale bar: 50 μm

Supplementary Figure S6

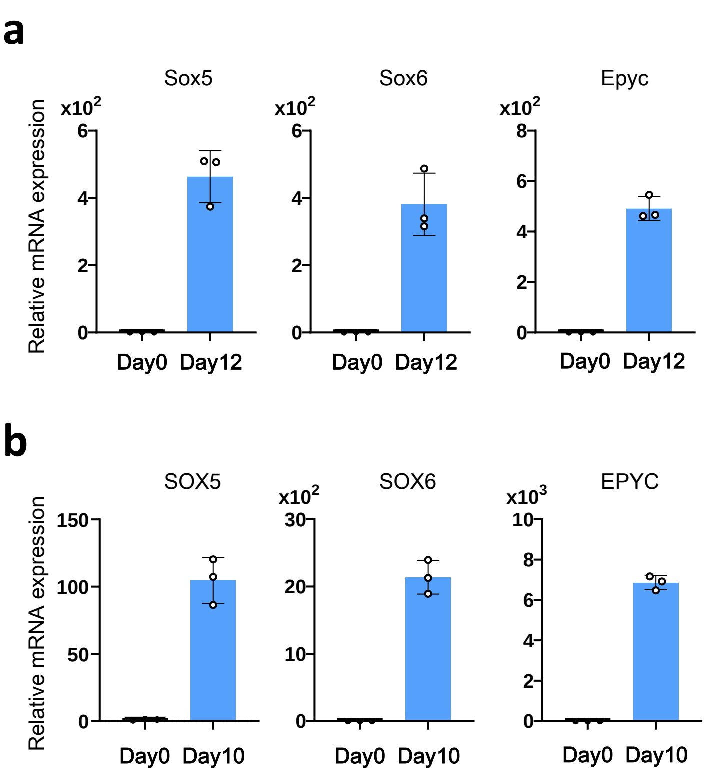


**Supplementary Figure S6 Expression of marker genes for chondrocyte differentiation in differentiating cells from mESC and hESC**

**a**, Results of qRT-PCR for *Sox5*, *6* and Epyc expression of mESC-derived trachea mesodermal cells at day0 and 12.

**b**, Results of qRT-PCR for *SOX5*, *6* and EPYC expression of hESC-derived trachea mesodermal cells at day0 and 10.

Each column shows the mean with S.D. (n=3). Scale bar; 50µm.

Source data are provided in Source data file.

Supplementary Figure S7

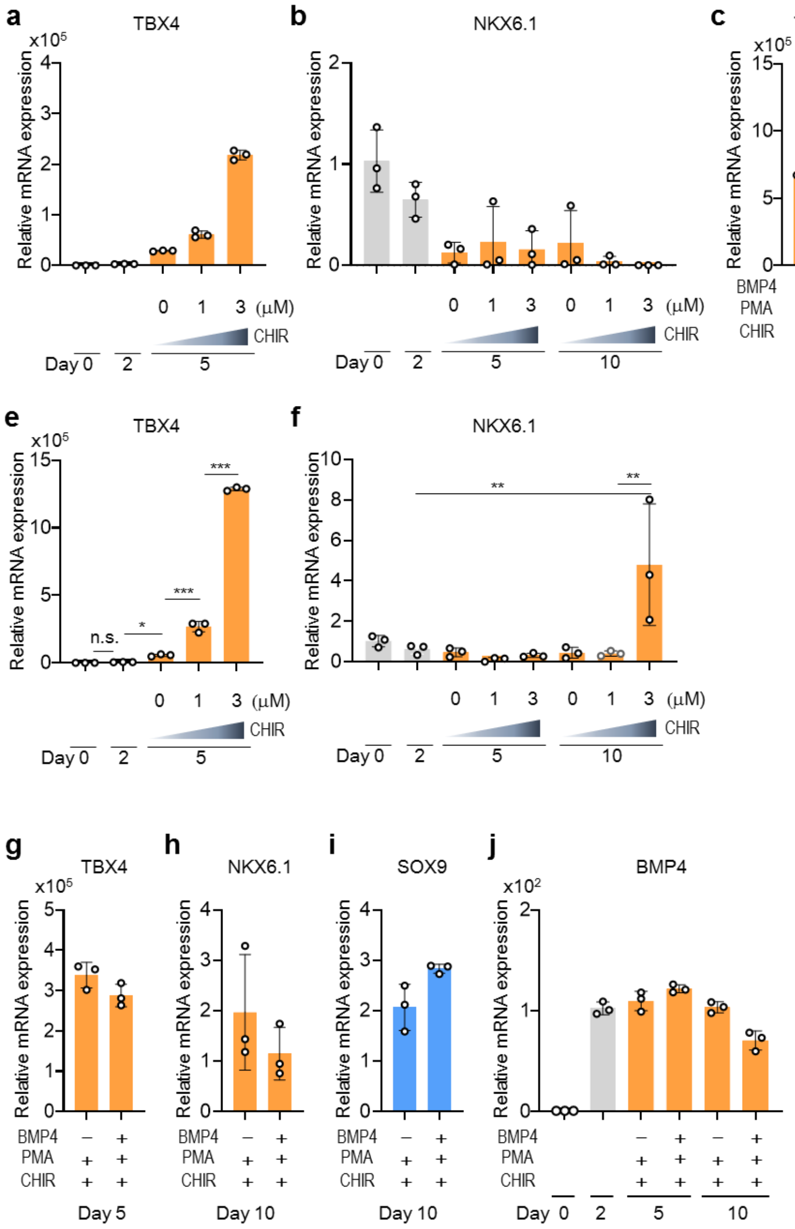


**Supplementary Figure S7 Indispensable role of the SHH agonist on trachea mesodermal differentiation from hESCs.**

**a** and **b**, Results of qRT-PCR for *TBX4* (**a**) and *NKX6.1* (**b**) expression in differentiating hESCs cultured with growth factor cocktails including BMP4 and different doses of Wnt agonist.

**c** and **d,** The results of qRT-PCR for *TBX4* (**c**) and *NKX6.1* (**d**) expression in differentiating hESCs cultured with growth factor cocktails including BMP4 and Wnt agonist with/without SHH agonist.

**e** and **f**, Results of qRT-PCR for *TBX4* (**e**) and *NKX6.1* (**f**) expression in differentiating hESCs. hESCs were incubated with growth factor cocktails including SHH agonist, BMP4 and different doses of Wnt agonist.

**g-i,** Results of qRT-PCR of *TBX4* (**e**), *NKX6.1* (**f**), *SOX9* (**g**) in differentiating hESCs cultured with/without BMP4.

**j**, Results of qRT-PCR for *BMP4* expression in differentiating hESCs at different time points. hESCs were cultured with/without BMP4.

Each column shows the mean with S.D. (n=3). *p<0.05, **p<0.005, ***p<0.0001

Source data are provided in Source data file.

### Supplementary Table 1 Methods for Immunohistochemistory

| *Antibody (Dilution)* | *Company, Catalog code* | *Fixative* | *Tissue preparation* | *Antigen retrieval* | *Secondary antibody* |
|---|---|---|---|---|---|
| *α-Smooth muscle actin(SMA)-Cy3 (1:1000)* | Sigma, C6198 | 4% PFA overnight at 4˚C | Paraffin | $Histo^{VT}$ One 15min at 105˚C | - |
| *GFP (1:200)* | Thermo Fisher Scientific, A10262 | 4% PFA overnight at 4˚C | Paraffin | $Histo^{VT}$ One 15min at 105˚C | Chicken Alexa488 |
| *Nkx2.1 (1:400)* | Santa Cruz Biotechnology, Inc, sc-13040 | 4% PFA overnight at 4˚C | Paraffin | $Histo^{VT}$ One 15min at 105˚C | Rabbit Alexa555 |
| *Nkx6.1 (1:100)* | Developmental Studies Hybridoma Bank, F55A12 | 4% PFA overnight at 4˚C | Paraffin | $Histo^{VT}$ One 15min at 105˚C | Mouse Alexa647 |
| *Sox2 (1:400)* | Santa Cruz Biotechnology, Inc, sc-17320 | 4% PFA overnight at 4˚C | Paraffin | $Histo^{VT}$ One 15min at 105˚C | Goat Alexa488 |
| *Sox9 (1:1000)* | Abcam, AB5535 | 4% PFA overnight at 4˚C | Paraffin | $Histo^{VT}$ One 15min at 105˚C | Rabbit Alexa488 |
| *Tbx4 (1:300)* | Abcam, ab220035 | 4% PFA overnight at 4˚C | Paraffin | $Histo^{VT}$ One 15min at 105˚C | Rabbit Alexa594 |

### Supplementary Table 2 Primer lists for the construction of *in situ* hybridization probes

| *Probe* | *Primer sequence (Forward, 5' to 3')* | *Primer sequence (Reverse, 5' to 3')* | *Accession number* | *Nucleotides* |
|---|---|---|---|---|
| *Wnt2* | ATAGTCGACACAGAGATCACAGCCTCTTT | ATAGAATTCCATGTCCTCAGAGTACAGGA | NM_023653 | 429-817 |
| *Wnt4* | ATAGTCGACAGAAACTCAAAGGCCTGATC | ATAGAATTCTCGTTGTTGTGAAGATTCATGA | NM_009523 | 176-67 |
| *Wnt5a* | ATAGTCGACATGGCTTTGGCCACGTTT TT | ATAGAATTCATTTGCATCACCCTGCCAAA | NM_001256224 | 95-402 |
| *Wnt7b* | ATAGTCGACTGAACCTTCACAACAATGAGG | ATAGAATTCAGTGGAATTTGCAGTTACACT | NM_001163633 | 947-1396 |

### Supplementary Table 3 Methods for RNAscope experiments

| *Probe* | *Catalog code* | *Fixative* | *Tissue preparation* | *Target Retrieval* | *Protease* | *Fluorophore* |
|---|---|---|---|---|---|---|
| *Axin2* | 400331-C2 | 4% PFA overnight at 4˚C | Frozen | 1xTarget retrieval 5min at 98-102˚C | Protease III 15min at 40 ˚C | Opal520 |
| *Nkx2.1* | 434728-C3 | 4% PFA overnight at 4˚C | Frozen | 1xTarget retrieval 5min at 98-102˚C | Protease III 15min at 40 ˚C | Opal570 |
| *Tbx4* | 483291 | 4% PFA overnight at 4˚C | Frozen | 1xTarget retrieval 5min at 98-102˚C | Protease III 15min at 40 ˚C | Opal690 |

### Supplementary Table 4 Methods for Immunocytochemistory

| *Antibody (Dilution)* | *Company, Catalog code* | *Fixative* | *Secondary antibody* |
|---|---|---|---|
| *Aggrecan (1:100)* | Abcam, ab3778 | 4% PFA 30 min at RT | Mouse Alexa647 |
| *α-SMA (1:200)* | Sigma, #A2547 | 4% PFA 30 min at RT | Mouse Alexa647 |
| *α-SMA-Cy3 (1:1000)* | Sigma, C6198 | 4% PFA 30 min at RT | - |
| *Collagen1a1 (1:200)* | Abcam. ab34710 | 4% PFA 30 min at RT | Rabbit Alexa546 |
| *Collagen2a1 (1:200)* | Santa Cruz Biotechnology, Inc, sc-7764 | 4% PFA 30 min at RT | Goat Alexa488 |
| *Foxf1 (1:500)* | R&D systems, AF4798 | 4% PFA 30 min at RT | Goat Alexa488 |
| *Gata4 (1:200)* | Santa Cruz Biotechnology, Inc, sc-1237 | 4% PFA 30 min at RT | Goat Alexa488 |
| *Nkx6.1 (1::100)* | R&D systems, AF5857 | 4% PFA 30 min at RT | Goat Alexa594 |
| *Sox9 (1:1000)* | Abcam, AB5535 | 4% PFA 30 min at RT | Rabbit Alexa488 |
| *Tagln (1:200)* | Abcam, ab14106 | 4% PFA 30 min at RT | Rabbit Alexa 594 |

Supplementary Table 5 Primer lists for quantitative RT-PCR (Mouse)

| | *Primer sequence (Forward, 5' to 3')* | *Primer sequence (Reverse, 5' to 3')* |
|---|---|---|
| *Acta2* | ACTGGGACGACATGGAAAAG | GTTCAGTGGTGCCTCTGTCA |
| *Bmp4* | GCCGAGCCAACACTGTGAGGA | GATGCTGCTGAGGTTGAAGAGG |
| *Epyc* | TTCTGGGTCCACACACCAAC | CTTCTTGGGCAGTGGAGGAATA |
| *Foxf1* | CCTGTCTGGCAGCATCTCCAC | GACTGTGAGTGATACCGAGGGA |
| *Gapdh* | TCACCACCATGGAGAAGGC | GCTAAGCAGTTGGTGGTGCA |
| *Gata4* | GCCTCTATCACAAGATGAACGGC | TACAGGCTCACCCTCGGCATTA |
| *Hoxb6* | GCTCTACTCGTCTGGCTATGC | GTGGGTAATAGGAGGACGCC |
| *Prrx1* | GACACCCCTCAGCAGGACAA | TGAAACCACACCTGCACTCT |
| *Sox5* | ACATGCACAATTCCAACATCAGC | GGTCATAGCTTTCCAGCGAGAT |
| *Sox6* | AATGCACAACAAACCTCACTCT | AGGTAGACGTATTTCGGAAGGA |
| *Sox9* | TGAGAGGTTTCAGATGCAGTG | CACATCCACATACAGTCCAGG |
| *Tbx4* | TCACTGGATGCGGCAGTTGGTCTCT | CACGTGGGTGCAAAAGGCTGTGTTT |
| *Tbx5* | GGACCCAGTCCCTTGAATGG | TCCAGGCTGAGGAGTTCTAGGC |
| *Wnt2* | CCAACGAAAAATGACCTCGT | GGGAAGTCAAGTTGCACACA |

Supplementary Table 6 Primer lists for quantitative RT-PCR (Human)

| | *Primer sequence (Forward, 5' to 3')* | *Primer sequence (Reverse, 5' to 3')* |
|---|---|---|
| *ACTA2* | CTATGCCTCTGGACGCACAACT | CAGATCCAGACGCATGATGGCA |
| *BMP4* | CAAACTTGCTGGAAAGGCTC | CCGCTACTGCAGGGACCTAT |
| *EPYC* | AGGAGGAGGAATCTACTCCCA | CAGCGGAGGAATAGCATCAAG |
| *FOXF1* | AGCAGCCGTATCTGCACCAGAA | CTCCTTTCGGTCACACATGCTG |
| *GAPDH* | CCCATCACCATCTTCCAGGAG | CTTCTCCATGGTGGTGAAGACG |
| *GATA4* | TAGCCCCACAGTTGACACAC | GTCCTGCACAGCCTGCC |
| *HOXB6* | CACTCCGGTCTACCCGTGGATGCA | CATATCTTGATCTGCCTCTCCGTCAG |
| *NKX6.1* | ATGACAGAGAGTCAGGTCAAGG | CTCCGAGTCCTGCTTCTTCTT |
| *PRRX1* | TGCAGGCTTTGGAGCGTGTCTT | CTCATTCCTGCGGAACTTGGCT |
| *SOX5* | CAGCCAGAGTTAGCACAATAGG | CTGTTGTTCCCGTCGGAGTT |
| *SOX6* | TTACTCGGCCAGAAGATGCAG | ACTCGTGCTTCAGCCACAGT |
| *SOX9* | GTAATCCGGGTGGTCCTTCT | GTACCCGCACTTGCACAAC |
| *TBX4* | TGATCATCACTAAGGCTGGCAG | ACAGAACTTGTAGCGATGGTCAT |
| *TBX5* | ACAAAGTGAAGGTGACGGGCCTTA | ATCTGTGATCGTCGGCAGGTACAA |
| *WNT2* | CTGTATCAGGGACCGAGAGG | CCCACAGCACATGACTTCAC |
